# Alginate-εPLL core-shell hydrogel beads as a tool for an effective, stable and scalable microbial encapsulation

**DOI:** 10.1101/2025.01.17.633598

**Authors:** Alba Amaro-Cruz, Miguel García-Román, Ignacio Moya-Ramírez

## Abstract

Synthetic microbial consortia have a significant potential for novel biotechnological applications such as the upgrade of complex feedstocks. However, achieving stable and reproducible co-cultures is challenging due to competitive dynamics and unbalanced growth rates among species. Here, we present an effective method for microbial encapsulation relying on alginate core-shell hydrogel beads coated with ε-poly-L-lysine (εPLL-HB). This procedure ensures complete containment of several microbial species (prokaryotic and eukaryotic) while allowing for consistent growth inside the beads. Compared to chitosan and α-poly-L-lysine, two of the biomaterials most used as coating agents for this kind of encapsulation, εPLL demonstrated superior performance in avoiding cellular escape across diverse culture conditions and for all the microbial strains tested. εPLL-HB enabled the construction of spatially distributed co-cultures, effectively balancing populations between microorganisms with different growth rates. Furthermore, microbial encapsulation inside εPLL-HB provided protection against toxic compounds in lignocellulosic-derived media and maintained the encapsulation efficacy and cell viability after long-term storage at −80 °C. The superior microbial containment, structural integrity and chemical resistance of εPLL-HB, combined with their cost-effective and simple preparation, makes them a versatile tool for synthetic microbial consortia engineering, with broad applicability in biotechnological processes.

## 1. Introduction

Communities of co-occurring and interacting microorganisms are complex systems ubiquitous in nature and represent the natural endpoint in most environments where microorganisms have naturally evolved (Sanchez et al., 2023; Zhang et al., 2021). Microbial consortia have the potential to challenge the current single strain-based paradigm in microbial biotechnology (Kost et al., 2023). Examples of these traits are the enhanced metabolic capabilities due to division of labour, which can result in reduction of metabolic burden and higher productivities or wider range of metabolizable carbon sources compared to single-species cultures (Lin, 2022; M. Wang et al., 2022). They also show better stability and resilience to changes in the environment (Liu, 2023). Natural microbial consortia have been traditionally used for composting, anaerobic digestions or wastewater treatment (Zhou et al., 2024), while synthetic ones, normally referred as co-cultures, have been developed with several purposes such as synthesis of biochemicals or environmental remediation, for example (Hu et al., 2017; Jiang, 2020; Millán Acosta et al., 2021; Senne de Oliveira Lino et al., 2021; Shahab et al., 2020; Sulbaran-Bracho et al., 2023).

However, harnessing the full potential of microbial consortia requires effective strategies to manipulate and engineer them (Sánchez et al., 2024). Engineering synthetic microbial consortia requires mechanisms to maintain the balance among co-cultured species (L. Wang et al., 2022). In liquid microbial co-cultures, differences in the growth and nutrient consumption dynamics between the species and other competitive interactions will normally lead to one species outgrowing and displacing the rest. This lack of populations control dramatically reduces the number of the microorganisms that can be co-cultured (Moya-Ramírez et al., 2022). Addressing this limitation is crucial to obtain stable and reproducible synthetic microbial consortia, thus making them suitable for their engineering and scaling-up and widely applicable to fields such as the valorisation of complex feedstocks like those derived from lignocellulosic side products.

In this regard, spatial distribution and microbial encapsulation allow a precise control over microbial populations in co-cultures (Johnston et al., 2020; Moya-Ramírez et al., 2022). Besides that, microbial encapsulation offers additional advantages such as mimicking microbial biofilms-based niches or protection against environmental stressors and toxic substances (Jeong and Irudayaraj, 2023). Among the available options for cell encapsulation, hydrogel-based technologies stand out for their biocompatibility, tuneable properties and ease of fabrication. These biomaterials also allow the mass transfer of biomolecules, maintaining the chemical communication with the exterior needed for nutrient uptake or cell signalling for instance. More specifically, alginate hydrogels excel on all these features, and therefore, have been extensively studied for cell encapsulation (Lee and Mooney, 2001; Paredes Juárez et al., 2014). However, the physical integrity of ionotropic hydrogels like alginate is highly sensible to chelating agents such as citrate or phosphate, and to cations such as Na^2+^ or Mg^2+^, that are commonly present in most culture media (Kim et al., 2014; Simó et al., 2017). Additionally, the growth of the encapsulated microbes or osmotic pressures can cause swelling of the alginate hydrogel (Jeong and Irudayaraj, 2023; Simó et al., 2017), which could lead to the loss of its structural integrity. For this reason, alginate-based core-shell hydrogels, where alginate beads containing the encapsulated cells are coated with other material that improves the mechanical and chemical resistance of alginate are a common approach used in this field (Kim et al., 2014; Li et al., 2017; Simó et al., 2017; Tang et al., 2021).

When applied to microbial co-cultures, avoiding cell escape from their confined space is another desired feature in any biomaterial used for cell encapsulation. Otherwise, the population balance could be compromised by the uncontrolled growth of the released microbes. Despite of the wide range of possible coating materials and techniques described for alginate core-shell microbial encapsulation, few of them address cell leakage. To our knowledge, and only recently, Wang et al. were able to co-culture several bacterial and yeast strains based on alginate and chitosan-coated beads, that by a preliminary analysis did not show cell escape (L. Wang et al., 2022) and Jeong et al. reported core-shell alginate capsules, with coating layers composed however exclusively of alginate, and systematically evaluated the conditions where no cell escape could be detected (Jeong and Irudayaraj, 2023).

In the present work, we aim to develop a microbial encapsulation system focused on ensuring a total containment of the encapsulated cells and their descendants. We describe a methodology suitable for encapsulating a wide variety of microbial species (prokaryotic and eukaryotic) that ensures both total microbial entrapment and their growth inside the hydrogel beads (HB) in standard culture conditions. The efficacy of the encapsulation is maintained over incubation in diverse culture media, and during prolonged periods under storage at −80 °C. We have systematically evaluated the performance of several HB built using a layer-by-layer methodology, comparing three positively charged polymers as coating agents. Two of them are frequently used for this purpose: chitosan and α-poly-L-lysine (αPLL). Additionally, we included ε-poly-L-lysine (εPLL) in our study, which is less commonly used and turned out to be the best performing of them. Our methodology exclusively relies on natural, cost-effective and biodegradable coating polymers and does not require specialized equipment (Nie et al., 2021). Its simplicity and compatibility with microfluidics and other automated devices make it easily scalable and, therefore, expand the tools for creating synthetic microbial co-cultures based on alginate core-shell capsules.

## 2. Materials and methods

### 2.1. Microbial strains, culture conditions and chemicals

The bacterial and yeast strains used for this work were: *Pseudomonas putida* KT2440 and *Kluyveromyces marxianus* DSM 70792 purchased from the German collection of microorganisms and cell cultures (DSMZ, Germany); *Yarrowia lipolytica* CBS 7504 (W29) purchased from the Westerdijk fungal biodiversity institute (the Netherlands); *Escherichia coli* BL21 pIDMv5K-GREEN, constitutively expressing green fluorescent protein (GFP), provided by Sebastian S. Cocioba from Binomica Labs and *Bacillus subtilis* B5 isolated from oil-rich environments which belongs to our strains collection. Stock cultures of microbial strains were maintained at −80 °C in LB (bacterial strains), LB with 50 μg/mL of kanamycin (*E. coli* BL21 pIDMv5K-GREEN) or GPY (yeast strains) supplemented with 25% (v/v) glycerol. Strains were streaked on LB or GPY agar plates and incubated at 30 °C for 24 h to obtain isolated colonies. *E. coli* BL21 pIDMv5K-GREEN was cultured in LB with 50 μg/mL of kanamycin at 37 °C.

Natamycin, kanamycin, citric acid, 2-[4-(2-hydroxyethyl)-1-piperazinyl]-ethanesulfonic acid (HEPES), EDTA, alginic acid sodium salt from brown algae (71238), low MW chitosan (ref: 448869), α-poly-L-lysine hydrochloride (αPLL, MW 15000–30000 Da, ref: P2658), M9 minimal salts and other inorganic reagents were supplied by MERCK (Germany). ε-poly-L-lysine hydrochloride (εPLL, MW 3500-4500 Da, ref: FP14985) was purchased from Biosynth (UK). LB and LB-agar media were obtained from VWR (USA). Yeast extract, peptone, D(+)-glucose and agar powder were supplied by Scharlab (Spain).

### 2.2. Culture media and buffers

The media used in this work were GPY, minimal medium (M9), and M9-coffee (M9c). GPY consisted of 40 g D(+)-glucose, 5 g peptone and 5 g yeast extract for 1L of solution. When necessary, 15 g/L agar powder was added to prepare GPY-agar. M9 was prepared with 33.7 mM Na_2_HPO_4_, 22.0 mM KH_2_PO_4_, 8.55 mM NaCl and 9.35 mM NH_4_Cl (M9 minimal salts), autoclaved and supplemented with 0.4% D(+)-glucose (w/v), 1 mM MgSO_4_, 0.3 mM CaCl_2_, 13.4 mM EDTA, and 3.1 mM FeCl_3_ from filter-sterilized stock solutions. M9c composition was 50% (v/v) M9 without D(+)-glucose and 50% (v/v) autoclaved spent coffee grounds hydrolysate. The detailed protocol for the obtention of this hydrolysate is described in the supplementary material. The buffers used were KRH, phosphate-buffered solution (PBS buffer), and disruption buffer. KRH buffer was prepared with 20 mM HEPES, 135 mM NaCl, 5 mM KCl, 0.4 mM K_2_HPO_4_, adjusted to pH 7.4, autoclaved, and supplemented with filter-sterilized 1 mM MgSO_4_ and 1 mM CaCl_2_. PBS buffer consisted of 137 mM NaCl, 2.7 mM KCl, 10 mM Na_2_HPO_4_, 1.8 mM KH_2_PO_4_ adjusted to pH 7.4 and autoclaved. Disruption buffer was prepared with 25 mM EDTA and 0.2 M citric acid adjusted to pH 7 and autoclaved. All autoclave sterilizations took place at 121 °C for 30 min, except for the GPY medium, which was sterilized at 115 °C for 15 min.

### 2.3. Encapsulation of microorganisms

A single colony of each microorganism was transferred into 5 mL LB or GPY in a sterile 50 mL Falcon^®^ tube (Corning) and incubated at 30 °C and 200 rpm for 9 h. These cultures were used as inoculum at 1% (v/v) of 5 mL of M9 medium and incubated overnight at the same conditions. Next, cells were pelleted at 9000 g for 5 min and resuspended in fresh M9 to an optical density (OD_600_) of 2. The microbial suspension was thoroughly mixed with a filter-sterilized 2% (w/v) sodium alginate solution (10 mM Tris pH 8.5) at a 1:3 volume ratio (0.25 and 0.75 mL, respectively) in an Eppendorf tube and drawn to a 1 mL syringe. The hydrogel beads (HB) were formed by dropping this mix through a sterile 30GA-blunt end needle (RS Components, U.K.) into 100 mL of 0.1 M CaCl_2_ solution from a height of 1 cm. Alginate HB were crosslinked for 30 min under magnetic agitation at 220 rpm. Then, they were collected in a sterile cell strainer (VWR, USA), washed with fresh CaCl_2_ solution, and incubated 1 min in KRH buffer. In this way we obtained the microbe-loaded HB cores used for subsequent experiments.

The layer-by-layer coating technique was employed to sequentially add layers of coating polymers onto the HB core, resulting in core-shell HB with varying number of coating layers (Figure 1). The method was based on the one described by Moya-Ramírez et al (Moya-Ramírez et al., 2022). Three different polymers were tested as coating agents: low MW chitosan, α-poly-L-lysine hydrochloride (αPLL) and ε-poly-L-lysine hydrochloride (εPLL). To this end, HB cores were transferred to a 15 mL Falcon^®^ tube containing 5 mL of 1 mg/mL solution of polylysine (either εPLL or αPLL) on KRH buffer, incubated for 10 min under gentle agitation in an oscillating shaker and washed with fresh KRH buffer. The HB were added straightaway to 80 mL of 0.2% (w/v) alginate solution (10 mM Tris pH 8.5) and kept under magnetic stirring at 220 rpm for 10 min. Next, the HB were rinsed with KRH buffer and incubated again with εPLL/αPLL-KRH solution for 10 min, washed with 0.15 M sterile mannitol, transferred to 5 mL of 1 mg/mL solution of εPLL/αPLL in 0.15 M mannitol and incubated for 2 h. The resulting beads will be referred onwards as three-layered εPLL or αPLL hydrogel beads (3L-εPLL-HB or 3L-αPLL-HB respectively). Finally, HB with an additional outer layer of alginate were also produced. For that, 3L-HB were rinsed with KRH buffer and incubated in 0.2% (w/v) alginate solution for 10 min, washed with KRH buffer and submerged in a crosslinking solution with 50 mM BaCl_2_ and 0.15 M mannitol for 5 min under magnetic agitation at 220 rpm. These HB will be referred as four-layered HB (4L-εPLL-HB or 4L-αPLL-HB). After the addition of the last coating layer, HB were rinsed and kept in PBS buffer. The protocol described was modified when chitosan was used as coating polymer, as detailed in the supplementary material.

**Figure 1.**
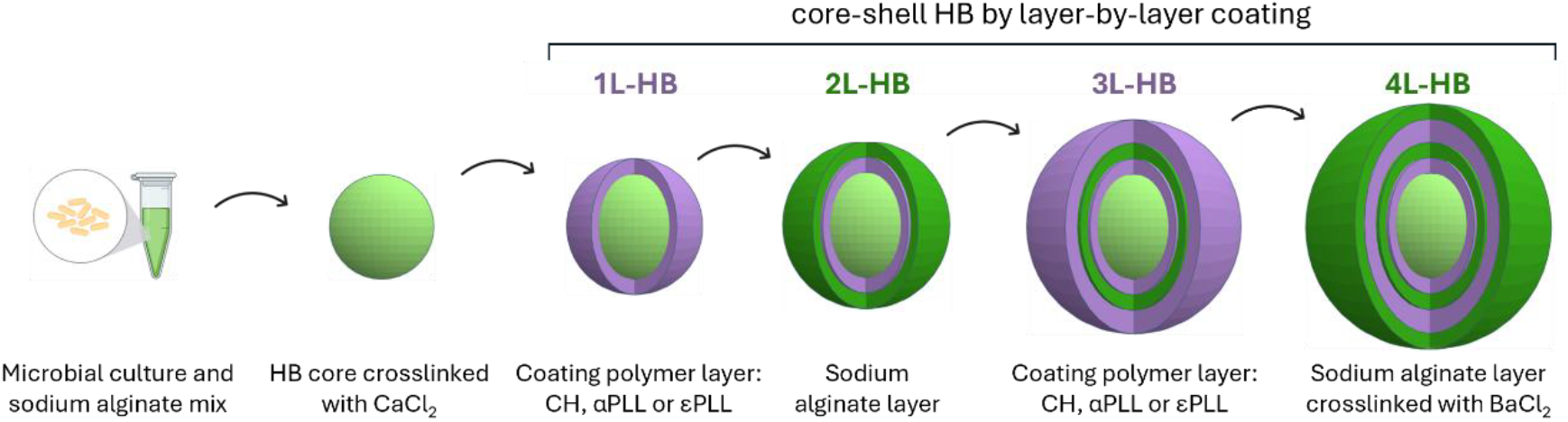
Schematic representation of the encapsulation and layer-by-layer coating method to obtain microorganism-loaded HB with three (3L-HB) and four coating layers (4L-HB).

### 2.4. Encapsulation efficacy, microbial viability and growth inside HB

Microbe-loaded HB were incubated in culture media to evaluate microbial survival and encapsulation efficacy. A single HB was placed in a well of a 48-well plate (Costar^®^) containing 0.5 mL of M9, LB or GPY medium. The HB was incubated at 30 °C and 150 rpm for 48 h. Viability was assessed by crushing the HB and plating its content on agar plates. The efficacy of the encapsulation was evaluated by the presence or absence of microorganisms in the liquid culture medium, assessed by plating it on agar medium. The encapsulation was considered as effective if no colonies were detected on them. HB prepared with chitosan, αPLL or εPLL as coating agents and with 3 and 4 layers of coating were tested. Wells without HB served as controls, and each condition was replicated at least eight times.

We also evaluated the growth of each strain encapsulated in HB. For that, just-prepared HB (0 h) and HB incubated for 24 and 48 h, were transferred to a sterile Eppendorf tube containing 1 mL of disruption buffer and vortexed for 10 min to detach the coatings of the capsules and enable the disruption of the alginate core. Homogenization was achieved by manually crushing the HB using a pipette tip and pipetting it until the complete homogenization of the core. Colony forming units (CFU) were then determined by plating serial dilutions of the cell suspensions on LB or GPY agar plates for bacteria and yeast respectively.

### 2.5. HB characterization by microscopy

HB shape, size and microbial distribution were examined using an Olympus CKX53 inverted microscope and a Leica DM IL LED fluorescence microscope (Leica Microsystems, Germany, excitation/emission: 390/450 nm). Methylene blue-stained and unstained HB were visualized immediately after preparation and after incubation in M9, LB or GPY media for 24 and 48 h. Methylene blue-stained HB were obtained by mixing the HB with a 0.01% methylene blue solution for 2 min under gentle agitation. The HB was then washed twice with distilled water.

Scanning electron microscopy (SEM) was used to examine the surface and cross-sectional structure of HB. Prior to observation, HB were fixed in 2.5% glutaraldehyde (0.05 M sodium cacodylate buffer) for 24 h at 4 °C. After the removal of glutaraldehyde with 0.1 M sodium cacodylate buffer, post-fixation was performed with 1% osmium tetroxide for 1 h at room temperature. Samples were then dehydrated through a graded ethanol series, reaching 100% ethanol, and subsequently subjected to critical point drying with CO2 using a Leica EM CPD300 critical point dryer. Finally, HB were cut in half with a razor blade and coated with carbon using an Emitech K975X coater. After processing, HB were observed under a high-resolution field emission SEM AURIGA (Carl Zeiss SNT) equipped with an AZtec detector (Oxford Instruments).

### 2.6. Spatially distributed co-cultures using HB

Spatially distributed co-cultures (SDCC) combining encapsulated *P. putida* inside 3L-εPLL-HB and planktonic *K. marxianus* cells were performed. These consisted of 20 HB in 100 mL Erlenmeyer flasks containing 10 mL of M9-LB medium (90% v/v M9 + 10% v/v LB), inoculated with 3.67 × 10^5^ CFU. In this way, the planktonic cells seed density was equivalent to the CFU inside of the 20 just-prepared *P. putida* 3L-εPLL-HB. Flasks were incubated at 30 °C and 250 rpm for 24 h. The growth of both microbes in the SDCC was compared to suspension co-cultures of both strains and their respective monoculture in the same conditions. Microbial growth was determined by plate counting. *P. putida* cell count was performed on LB plates supplemented with 20 μg/mL of natamycin and *K. marxianus* on GPY plates supplemented with 50 μg/mL of kanamycin. All experiments were conducted in duplicate.

### 2.7. Evaluation of cell viability during HB storage

Just-prepared 3L-εPLL-HB containing *B. subtilis* were incubated for 1 h in storage solution, consisting of M9 medium supplemented with 5% (w/v) mannitol under stirring at room temperature. HB were aliquoted and stored in fresh storage solution. Refrigeration at 4 °C and freezing at −80 °C were tested as storage conditions. To assess the effect of the storage on *B. subtilis*-loaded HB, encapsulation efficacy, microbial viability and growth in M9 medium for 24 h were evaluated as described in section 2.4. Stored HB were also characterized under inverted microscope.

### 2.8. Toxicity tolerance test of microbe-loaded HB

The ability to grow in lignocellulosic byproducts and the tolerance of toxic compounds on it were evaluated for planktonic and encapsulated cells cultured in M9c. For that, suspension cultures at different seed density were compared to cultures of the same HB-encapsulated microbe varying the number of HB in the well (from one to five). All cultures were performed in 48-well plates and incubated in a CLARIOstar Plus Microplate Reader (BMG LABTECH) at 30 °C and 300 rpm for 48 h. OD_600_ of suspension cultures was read every 20 min. Following incubation, cell viability and encapsulation efficacy were evaluated as explained in section 2.4.

## 3. Results

### 3.1. Encapsulation efficacy of the different coating agents

The most important feature of the encapsulation method that we aim to develop is to ensure complete microbial containment. Therefore, we decided to optimize the coating agent used to build core-shell alginate hydrogel beads (HB). We first encapsulated several microbial strains (prokaryotic and eukaryotic) in alginate hydrogels using an extrusion-dripping method. Following the gelation of the alginate microbe-loaded HB cores, we applied a layer-by-layer coating technique to add successive intercalate layers of coating polymers (chitosan, αPLL or εPLL) and alginate (Figure 1).

An antimicrobial assay showed that chitosan did not inhibit microbial growth (Table S1 and Figure S1). Conversely, αPLL and εPLL, which are commonly used as antimicrobial agents (Lebaudy et al., 2023; Ye et al., 2013; Zhu et al., 2023), inhibited the growth to varying degrees depending on the strain. However, this inhibitory effect did not hinder their use as coating agents, since all strains tested remained viable after their encapsulation and incubation regardless the coating polymer and the number of coating layers.

Next, we evaluated the encapsulation efficacy by incubating the polymer-coated HB in M9 medium for 48 h. Preliminary tests showed that HB coated with one and two layers rapidly lost their stability and disintegrated, so we focused on evaluating the biocontainment of HB with three and four layers of coating. Table 1 summarizes the results obtained. Three-layered chitosan-HB (3L-CH-HB) failed to entrap any of the microorganisms; however, the additional alginate outer layer of 4L-CH-HB ensured the containment of *B. subtilis* and *K. marxianus*. αPLL-HB performed better and successfully encapsulated *B. subtilis* and *K. marxianus*, regardless of the coating layers in the HB. However, *P. putida* escaped from both 3L and 4L-αPLL-HB, and *Y. lipolytica* remained entrapped in αPLL-HB only during 24 h of incubation. In contrast, εPLL-HB successfully encapsulated all bacterial and yeasts strains, with both the 3L and 4L coatings during 48 h of incubation. Next, we replicated the same experiment in rich media, LB or GPY for bacteria and yeast respectively (Table S2). Under these conditions, *B. subtilis* and *K. marxianus* encapsulated in 4L-CH-HB and in 3L and 4L-αPLL-HB escaped from the beads. Remarkably, both 3L and 4L-εPLL-HB retained their ability to contain the four strains when incubated in rich media during 48 h.

**Table 1.**
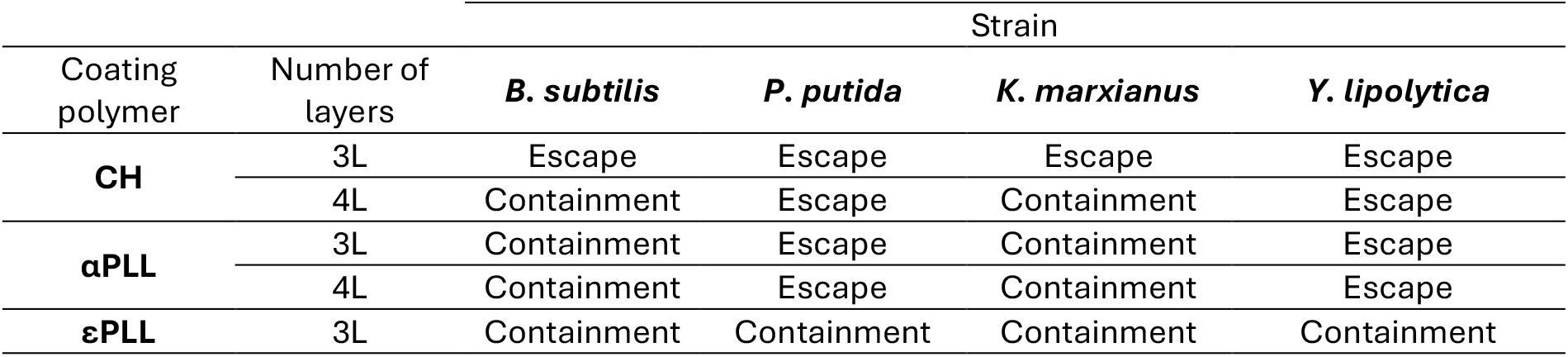
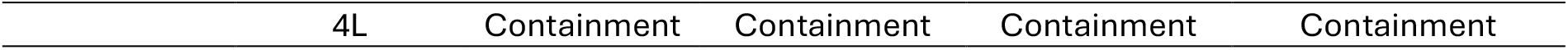
Evaluation of the encapsulation efficacy of each microbial strain in CH-, αPLL-, and εPLL-HB with three (3L) or four (4L) coating layers. HB were cultured in 0.5 mL of M9 medium for 48 h in individual wells.

| Coating polymer | Number of layers | Strain |  |  |  |
| --- | --- | --- | --- | --- | --- |
|  |  | *B. subtilis* | *P. putida* | *K. marxianus* | *Y. lipolytica* |
| CH | 3L | Escape | Escape | Escape | Escape |
|  | 4L | Containment | Escape | Containment | Escape |
| αPLL | 3L | Containment | Escape | Containment | Escape |
|  | 4L | Containment | Escape | Containment | Escape |
| εPLL | 3L | Containment | Containment | Containment | Containment |

Based on these results, we selected εPLL as the best performing coating agent and focused on its characterization and applications.

### 3.2. Microbial growth inside εPLL-HB

To assess whether HB can sustain microbial growth, we encapsulated the four microbial strains in εPLL-HB and cultured them in M9 and rich medium (LB for bacteria and GPY for yeasts). We initially observed their shape, size and structure. Control εPLL-HB (not loaded with microorganism) with three and four coating layers displayed a spherical shape after their preparation, with dimeter of the alginate core of 1944 ± 24 μm and a shell thickness of 141± 7 μm for a 3L-HB and 194 ± 8 μm for a 4L-HB (Table S3).

Next, we analysed the growth and spatial distribution of the microbial cells within the HB by optical microscopy. As showed in Figure 2a, the distribution varied depending on the encapsulated strain, the culture medium, and the number of coating layers in the HB. In M9 medium, bacterial and yeast strains in 3L-εPLL-HB spread throughout most of the alginate core. The addition of an extra coating layer led to the formation of dense colonies aggregates of *B. subtilis* and *P. putida*, concentrated in the central region of the alginate core. Similarly, *Y. lipolytica* exhibited greater density in the central region but it also grew towards the limits of the HB. In contrast, *K. marxianus* displayed a uniformed growth pattern in 4L-εPLL-HB. In rich medium, bacterial strains in 3L-εPLL-HB showed a similar growth pattern to that observed in M9. However, in 4L-εPLL-HB, *B. subtilis* formed large clusters of colonies, while *P. putida* displayed a distinct aggregate morphology, with smaller colonies spreading throughout the entire alginate core with a higher density in the central region. In contrast, when culturing encapsulated yeasts in GPY medium, cells spread all over the HB core after 24 h (data not shown) and became mostly opaque after 48 h. It is worth noting that εPLL-HB surface wrinkled as a result of incubation, especially in rich media. However, they preserved their structural integrity and the strict biocontainment of the encapsulated microorganism.

**Figure 2.**
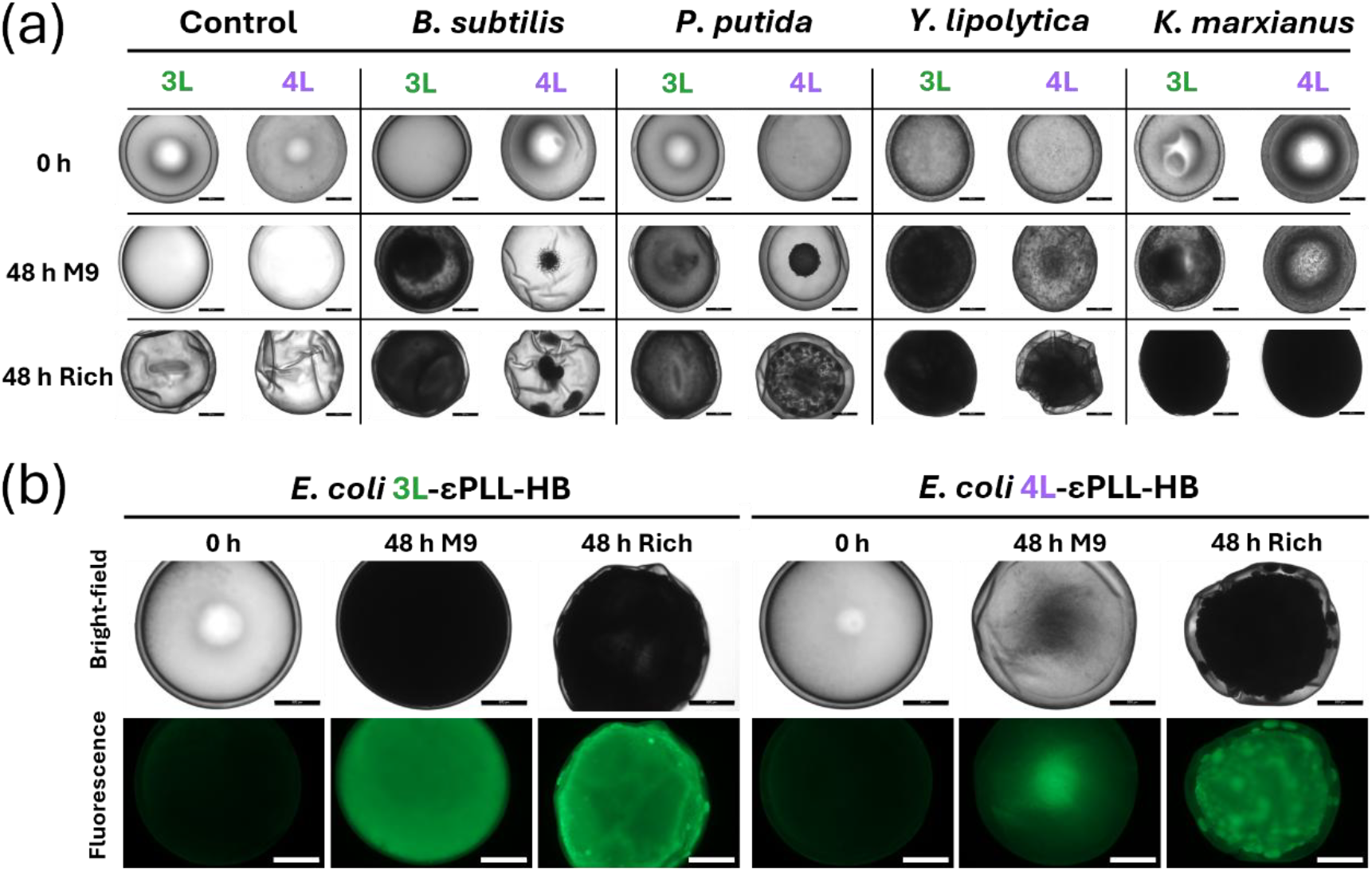
Bacterial and yeast strains encapsulated inside εPLL-HB with three (3L) and four layers (4L) of coating. **(a)** Optical microscopy images showing 3L and 4L-εPLL-HB not loaded with microorganism (control) and loaded with four different microbial strains. Images were taken after preparation (0 h) and after their incubation in M9 or rich medium (LB and GPY for HB containing bacteria and yeast strains, respectively) for 48 h. **(b)** Bright-field and fluorescence microscopy images showing 3L and 4L-εPLL-HB loaded with E. coli BL21 pIDMv5K-GREEN after preparation (0 h) and incubation in M9 or rich medium for 48 h. Scale bars: 500 μm.

To better elucidate the microbial distribution and growth patterns inside the HB, we encapsulated an *E. coli* strain constitutively expressing GFP. We clearly observed microbial growth after 24 h (data not shown) and 48 h, compared to just-prepared HB (0 h) (Figure 2b). Consistently with the previous optical microscopy images, the coating layers exhibited lower fluorescence intensity than the core, suggesting that the microbial proliferation is restricted to the HB core. These experiments also confirmed the influence of the culture medium and the number of coating layers on the growth pattern inside the HB.

After that, we studied the cell growth and density of the encapsulated microorganisms. For that, we disrupted the HB after 0, 24 and 48 h of incubation and determined the cell density inside the HB (CFU/mL, referred to HB volume). As shown in Figure 3, all strains were able to proliferate within εPLL-HB, showing a considerable increase in their CFU/mL after 24 h of incubation in both M9 and rich media. Notably, cell counts were higher in 3L-εPLL-HB compared to 4L, except for *Y. lipolytica*. As expected, CFU/mL were higher for HB incubated rich medium, in many cases reaching densities more than one order of magnitude higher compared to M9. These findings support the previous observations with optical microscopy. We also compared the cell density inside the HB with that in suspension cultures in Erlenmeyer flasks with rich medium (horizontal red lines in Figure 3). Our initial hypothesis was that microorganisms will reach lower cell densities when encapsulated inside HB. Indeed, this was the case for *P. putida* and *Y. lipolytica*. However, *B. subtilis* reached similar cell density growing in suspension in LB than inside 3L-εPLL-HB after 24 h and inside 4L-εPLL-HB after 48 h of incubation in this medium. Surprisingly, *K. marxianus* showed a markedly distinct behaviour to the rest of the encapsulated strains, since it reached remarkably high cell densities when encapsulated in both 3L and 4L-εPLL-HB, which were almost an order of magnitude higher compared to its planktonic cell density. This result refutes our initial hypothesis and shows that the behaviour of encapsulated microorganisms can significantly vary between different strains.

**Figure 3.**
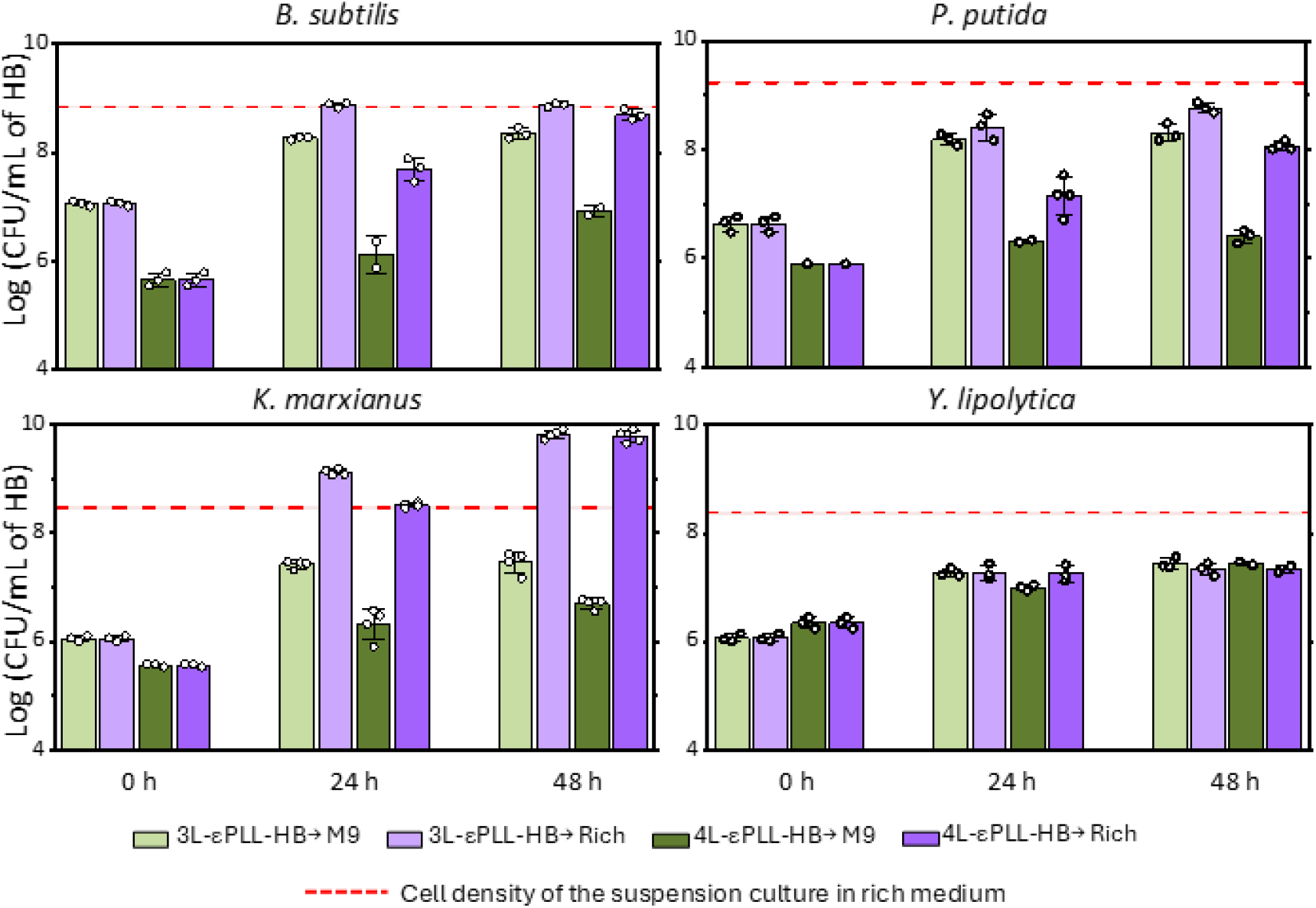
Log CFU per mL of εPLL-HB at different incubation times and culture media (M9 and rich medium, LB and GPY for bacteria and yeast respectively). At least two independent biological replicates were run (results for each experiment are indicated by circles), bars represent the average and error bars the standard deviation. The horizontal dashed red line represents the CFU/mL of the microorganism in rich medium growing in a suspension culture, and the shadowed area the SD.

### 3.3. εPLL-HB characterization by microscopy

To analyse the distribution and growth patterns of the encapsulated microorganisms in more detail, we examined stained εPLL-HB under optical inverted microscopy (Figure 4 and Figure S3). As previously mentioned, we observed different growth patterns depending on the encapsulated microorganism, the coating layers in the HB and the culture medium. Under higher magnification, we identified individual cells in just-prepared HB (0 h) located in both the core and the coating layers for the four encapsulated strains. However, only the cells located in the alginate core proliferated, as inferred from the lack of colonies in the coating layers.

**Figure 4.**
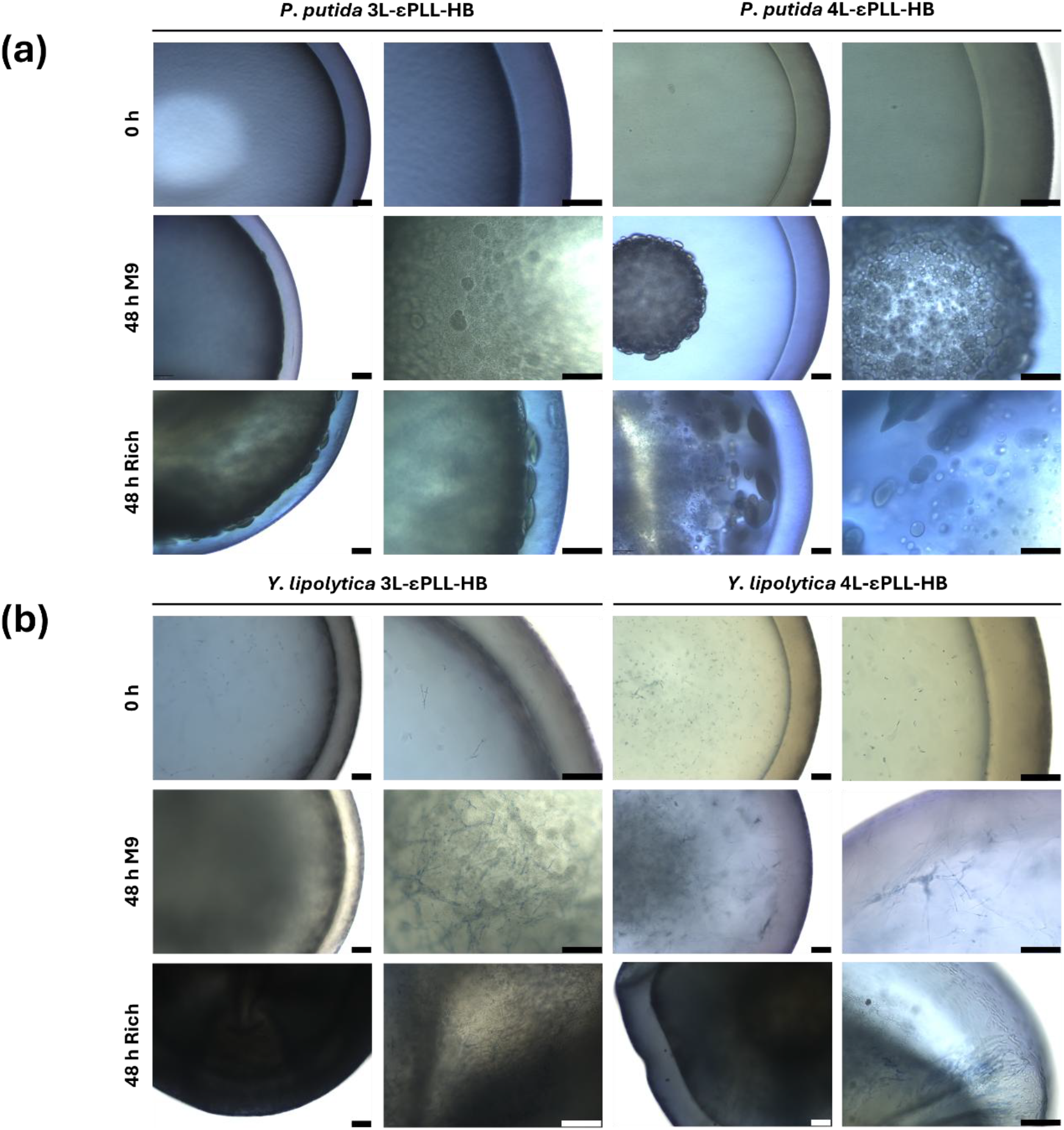
Microscopy images of methylene blue-stained εPLL-HB with three layers (3L) and 4 layers (4L) of coating loaded with *P. putida* **(a)** and *Y. lipolytica* **(b)**. The pictures show the just-prepared HB (0 h) and those incubated 48 h in M9 or rich medium (LB and GPY for HB containing bacteria and yeast strains respectively). Scale bars: 100 μm.

Bacterial strains encapsulated in 3L-εPLL-HB and incubated in M9 medium proliferated throughout the entire alginate core. The addition of a fourth coating layer led to the formation of large bacterial clusters of colonies concentrated in the central region of the HB core, with distinctive morphologies for each strain: *B. subtilis* grew forming clusters of colonies with an oval and irregular shape and size, whereas *P. putida* colonies also exhibited a flattened spheroid-like morphology. When incubating them in rich medium, colony aggregates protruded in some regions of the coating layers, particularly for *P. putida* in 3L-εPLL-HB, and they were denser, in agreement with the CFU counts described in section 3.2. Colonies in 4L-εPLL-HB were larger in size compared to those in 3L-εPLL-HB when incubated in rich medium for both strains, and in this case no inclusions were observed in the coating layer.

*K. marxianus* and *Y. lipolytica* are dimorphic yeasts known to undergo morphological changes in response to various factors. Microscopy images of just-prepared εPLL-HB revealed that *K. marxianus* exhibited a yeast-like morphology inside both 3L and 4L-HB, while *Y. lipolytica* displayed a pseudohyphal form. Consistent with our previous observations, yeasts strains colonized the entire core of 3L-HB after incubation in M9 medium. This pattern was also observed for *K. marxianus* encapsulated in 4L-HB, whereas the pseudohyphae of *Y. lipolytica* were primarily concentrated in the central region of the core, with some cells protruding into the coating layers. In rich medium, εPLL-HB containing *K. marxianus* became mostly opaque after 48 h of incubation in both 3L and 4L-εPLL-HB, limiting further observations. This is consistent with the high cell densities found for this yeast. In contrast, *Y. lipolytica* cells remained visible within the alginate core of both 3L and 4L-εPLL-HB spreading to the entire HB alginate core in the former and remaining concentrated in the centre in the latter.

Lastly, we observed the εPLL-HB using scanning electron microscopy (SEM). Cross-sectional images allowed us to clearly distinguish the core from the coating layers in both 3L and 4L-εPLL-HB controls with no cells inside (Figure S5). We observed significant structural differences depending on the number of coating layers. In 3L-εPLL-HB, the coating layers appeared separated from each other. In contrast, 4L-εPLL-HB exhibited a thicker and more compact alginate hydrogel core. As expected, the external structure of the HB varied depending on the outer coating polymer. In 3L-εPLL-HB the εPLL outer surface appeared smooth, whereas the addition of an alginate layer in 4L-εPLL-HB resulted in a rougher texture. SEM images of microbe-loaded εPLL-HB (Figure 4 and Figure S6) allowed us to confirm that the core-shell structure was preserved for both, 3L and 4L configurations, after 48 h of incubation in M9 medium. Interestingly, we found that the central region of 3L-εPLL-HB appeared almost empty as a consequence of microbial growth. On the other hand, the structure of the alginate core remained mostly unaltered in 4L-εPLL-HB, and groups of microbial cells were detected confined in specific regions of the alginate core, resembling the colonies observed in optical microscopy images. More importantly, while cells could be observed in contact with the inner region of the coating layer, no cells were detected in the outer side of the capsules. This corroborates the encapsulation efficacy of the method described.

### 3.4. Applications of the εPLL-HB

#### 3.4.1. Spatially distributed co-cultures

Following their characterization, we evaluated the applicability of εPLL-HB. For that, we employed them for the construction of a spatially distributed co-culture (SDCC) of two microbial strains, wherein one of them grew in suspension and the other encapsulated within 3L-εPLL-HB. For this purpose, we selected two strains with notable differences in their growth rate and cell density at their stationary phase: *P. putida* and *K. marxianus* (Figure 6a). When cultured in suspension in M9-LB medium, *K. marxianus* reached a density of 5.18 × 107 CFU/mL (Figure 6b.i), nearly two orders of magnitude lower than *P. putida*, and had a longer lag phase. However, in a suspension co-culture both strains inoculated at equal densities (3.67 × 10^5^ CFU/mL), *P. putida* reached a cell density comparable to its monoculture after 24 h, whereas *K. marxianus* experienced a drastic 50-fold reduction of its cell density (Figure 6b.ii). Conversely, when co-cultured as a SDCC, where *K. marxianus* grew in suspension and *P. putida* encapsulated within 3L-εPLL-HB, the former reached the same density than its monoculture, while the bacteria grew inside the εPLL-HB to the same extent than a control culture without *K. marxianus* in the liquid medium (Figure 6b.iii and iv).

**Figure 5.**
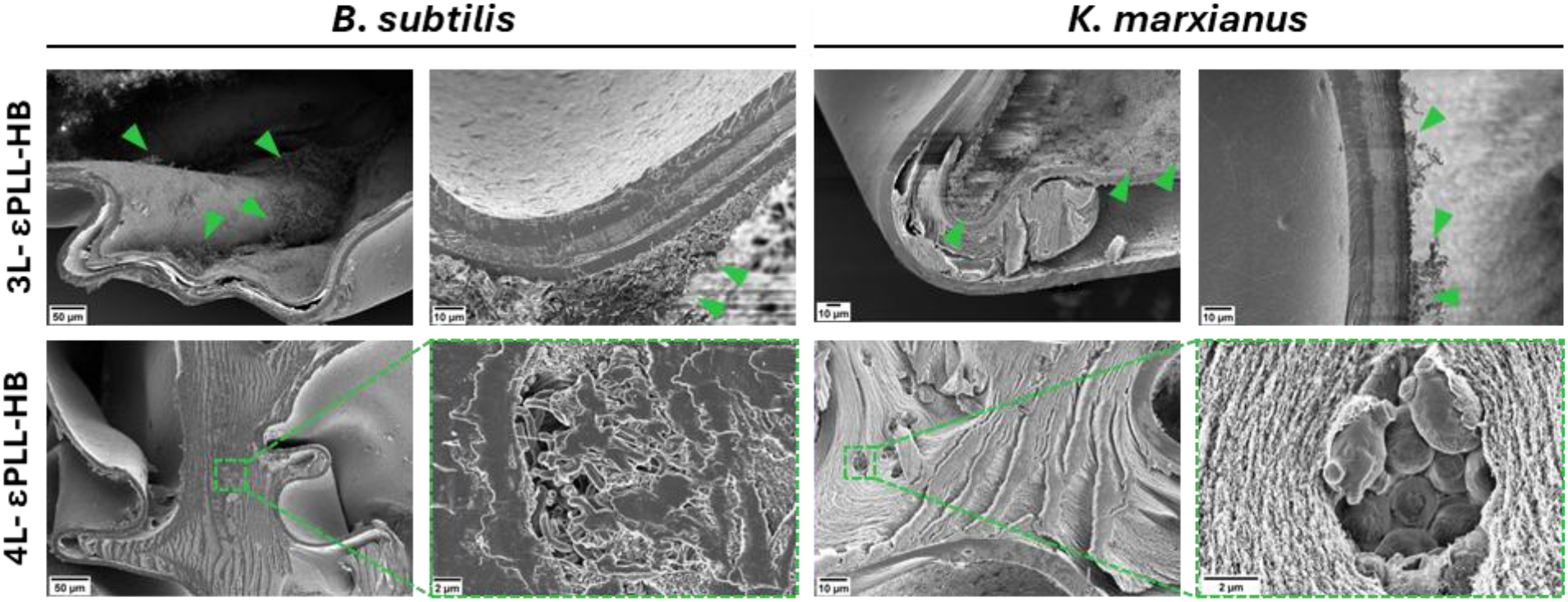
SEM images of εPLL-HB with three layers (3L) and 4 layers (4L) of coating loaded with *B. subtilis* and *K. marxianus* after their incubation for 48 h in M9 medium. Green triangles indicate microbial cells.

**Figure 6.**
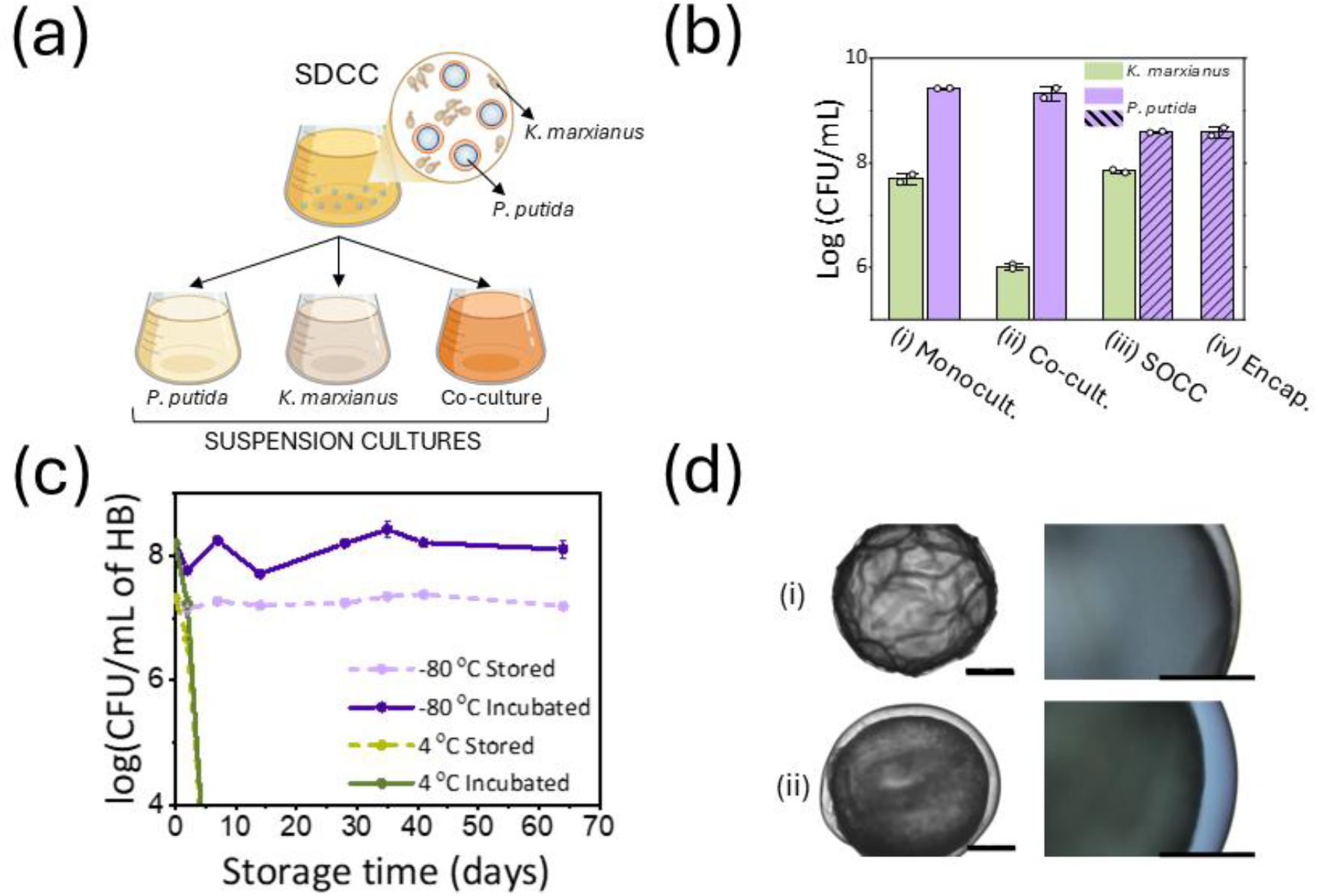
**(a)** Schematic representation of the spatially distributed co-culture (SDCC) configuration, consisting of encapsulated *P. putida* and planktonic *K. marxianus*. This system was compared with suspension monocultures of each strain and their co-culture. **(b)** Cell density of *K. marxianus* and *P. putida* grown in different conditions: (i) separately in suspension monocultures; (ii) combined in a suspension co-culture; (iii) as a SDCC with *K. marxianus* in suspension and *P. putida* encapsulated inside 3L εPLL-HB; and (iv) a control monoculture of *P. putida* growing inside εPLL-HB. Cell density units are expressed as log (CFU/mL), referred to the 0.5 mL of liquid medium for planktonic growing cells, and to εPLL-HB volume for encapsulated *P. putida*. Independent biological duplicates were run (results for each experiment are indicated by circles), bars represent the average and error bars the standard deviation. **(c)** Evolution of the cell density of *B. subtilis* encapsulated inside 3L-εPLL-HB and preserved at 4 °C and −80 °C. Cell density was measured after several storage times (up to 64 days) right after their recovery from their refrigerated or frozen storage and after their incubation in M9 medium during 24 h. Results correspond to the average of duplicates. **(d)** Optical microscopy images of HB stored at −80 °C after (i) thawing and (ii) 24 h of incubation in M9 medium. Scale bars: 500 μm.

#### 3.4.2. Long-term preservation of microorganism-loaded HB

We next tested the possibility to preserve 3L-εPLL-HB in storage solution under two conditions: refrigerated at 4 °C and frozen at −80 °C. We assessed the performance of 3L-εPLL-HB on retaining their ability to avoid any cell leakage while sustaining the growth of the encapsulated microorganism after storage compared to just prepared beads. To do so, we quantified the viable *B. subtilis* cells within 3L-εPLL-HB after their recovery from the refrigerated and frozen storage. Additionally, we incubated the 3L-εPLL-HB in M9 medium for 24 h and analysed the microbial growth inside of them (Figure 6c), as well as the culture medium to detect any cell escaping from the beads during the incubation.

The results reveal a big disparity between the two preservation conditions tested. At 4 °C, the cells inside 3L-εPLL-HB rapidly died, with a 10-fold decrease in viability after 2 days of preservation compared to just-prepared beads, and no detectable CFU after 7 days. In contrast, 3L-εPLL-HB frozen at −80 °C maintained the same viable cells as the just-prepared ones for at least 64 days. Accordingly, *B. subtilis* reached the same cell density inside 3L-εPLL-HB after their incubation in M9, regardless the duration of their storage at −80 °C. Notably, after the freezing and thawing process we observed HB forming ridges and grooves in their surface (Figure 6d). However, none of the tested 3L-εPLL-HB showed any cell leakage after their storage and subsequent incubation in M9 medium.

#### 3.4.3. Protection against harsh environmental conditions

We tested the four strains growing in suspension and inside 3L-εPLL-HB in M9-coffe (M9c) medium to explore the protective properties of the HB against the toxic and inhibitory compounds present in this lignocellulose-derived medium such as furfurals or phenolic compounds, which was produced through a hydrothermal hydrolysis pretreatment (Lara-Ramos et al., 2023; R. Wang et al., 2022). Figure 7a shows the influence of the seeding density over the growth kinetics for the four strains growing in suspension. The results varied between each strain. While *P. putida* seems to have a high susceptibility to the inhibitory and toxic effect of M9c and only showed a detectable OD_600_ increase when the seeding density was 10^7^ CFU/mL, *B. subtilis* performed better in this medium and grew for all the densities tested, albeit the lag phase was considerably extended when decreasing the seeding density. Both yeast strains performed similarly and only showed a significant growth with the two highest inoculum cell densities. Complementarily to the OD_600_ measurements, we sampled the cultures and plated a 20 μL drop in LB/GPY-agar in order to detect viable cells after the incubation time. None of the *P. putida* cultures had viable cells with the exception of the one that showed growth, which confirms that the bacteria died in all the cultures were an OD_600_ increase was not detected. In contrast, viable cells were found in all the *B. subtilis* cultures. Interestingly, for both yeasts, we also found viable CFU in those cultures where no OD_600_ increase was detected. Since the CFU found increased in number with their correspondent inoculum cell density, this observation suggests a sporulation induced by the M9c and that the yeasts did not replicate until they were seeded at a critical inoculum cell density.

**Figure 7.**
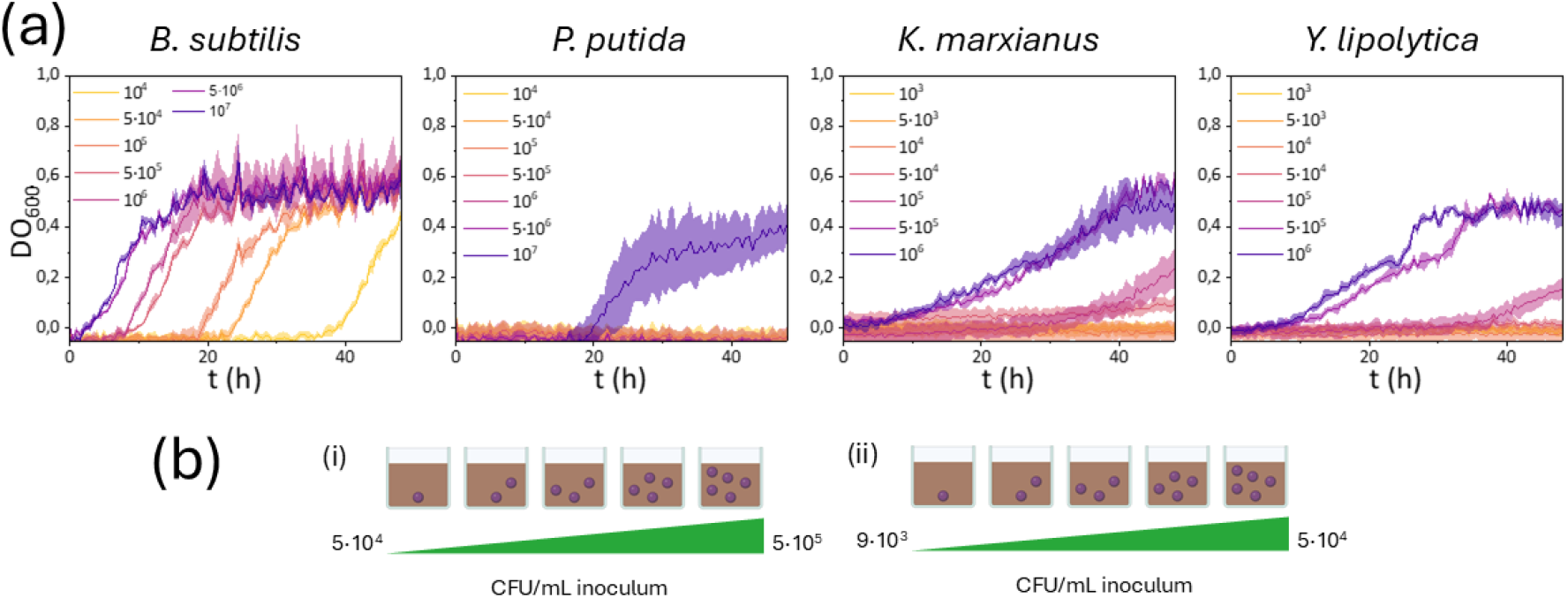
**(a)** Effect of the seeding density over the growth kinetics of liquid cultures of each of the strains tested. Cells were cultured in 0.5 mL of M9c medium at different seeding densities (ranging from 10^4^ to 10^7^ CFU/mL for bacteria and from 10^3^ to 10^6^ CFU/mL for yeast strains) in 24-well plates. The graphs show the OD600 blank subtracted using fresh M9c. Experiments were performed in triplicate; the straight line corresponds to the average OD600 and the shadowed area the standard deviation. **(b)** Schematic representation of the inoculum seed densities of bacteria (i) and yeasts (ii) encapsulated in 3L-εPLL-HB, as a function of the number of HB added per well (ranging from one to five).

We performed a similar experiment, in this case with encapsulated microbes, by adding between 1 and 5 εPLL-HB to 0.5 mL of M9c (Figure 7b). The equivalent seeding densities were calculated considering the total CFU per bead at the beginning of the experiment and the 0.5 mL of culture volume (detailed in Table S7). All of them were in the mid to lower range of those tested in the previous experiment. After 48 h of culture the beads were disrupted and their whole content plated on LB/GPY-agar. All the εPLL-HB kept viable CFU after their incubation on M9c. Interestingly, the cell density of the spread HB was similar, irrespective of the number of beads used, for all the strains except for *Y. lipolytica*. In this case, the CFU detected increased for experiments using one bead to those with five (Figure S8).

## 4. Discussion

### 4.1. Optimization of HB core coating agent

The main purpose of this work is to design a microbial encapsulation method that ensures complete microbial containment while allowing the encapsulated cells to proliferate inside them. To achieve that, the encapsulating material must be chemically and mechanically resistant to maintain its structural integrity and prevent cell escape during the culture. At the same time, the material should be flexible enough to allocate the cells as they multiply and maintain a chemical communication with the surrounding environment to allow them access to the nutrients needed for their growth. Considering all these factors, we selected alginate-based core-shell encapsulation as our approach.

We have tested the performance of three positively charged polymers as coating agents. Two of them, chitosan and αPLL, are commonly used to form core-shell alginate hydrogel beads (HB), while the third one, εPLL, has been scarcely explored to that purpose. In fact, to our knowledge, only Ma et al. reported the use of this polypeptide to obtain alginate capsules containing CHO cells (Ma et al., 2013). In all the tests run εPLL clearly outperformed the other two coating polymers. Regarding the structural integrity of the HB, both poly-L-lysines formed stable HB that maintained their structural integrity after their incubation in minimal medium, but chitosan-HB (CH-HB) only were stable when prepared with four coating layers (Figure S9a). We also observed some adhesion between CH-HB just after their preparation (Figure S9b), which can significantly hinder their applicability and compromise their encapsulation efficacy. Concerning the efficacy of the microbial containment inside the HB, we detected clear advantages of εPLL compared to αPLL: while no escape was detected from εPLL-HB, *P. putida* and *Y. lipolytica* escaped from αPLL-HB. Interestingly, *P. putida* is the smallest strain tested and *Y. lipolytica* was found to grow as pseudohyphae inside HB. Both, small size and pseudohyphae morphology can challenge the complete containment inside the beads, and only εPLL demonstrated to contain both microorganisms regardless of the coating layers. Additionally, under our experience, non-coated alginate beads and even some core-shell HB quickly destabilized in rich media like LB. However, it was not the case for HB coated with εPLL, which maintained their structural integrity and strict biocontainment of the encapsulated microorganism when incubated in rich media, as opposed to αPLL coated ones. Finally, swelling has been previously observed for beads formed only with alginate or chitosan-coated alginate during their incubation (Jeong and Irudayaraj, 2023; Moya-Ramírez et al., 2022). However, εPLL-HB did not show a significant swelling after their incubation (Table S4), which at last prevents the formation of cracks on the HB surface that could compromise its encapsulation efficacy.

Summarizing, εPLL showed a superior performance compared to the other two coating polymers tested. In addition, its price is orders of magnitude lower than αPLL, since it is naturally produced, while αPLL is a synthetic polymer (Ma et al., 2013). Therefore, we consider that εPLL is a promising coating material for the preparation of core-shell alginate HB, which outperformed two coatings agents commonly used by many cell encapsulation protocols.

### 4.2. Microbial growth inside εPLL-HB

Once εPLL was identified as the best performing coating agent, we evaluated its compatibility with the growth of diverse encapsulated microbes (prokaryotic and eukaryotic). The four tested strains multiplied by a factor of 10 to 1000 when cultured inside εPLL-HB, which demonstrates the compatibility of this polymer with the microbial growth in the alginate core of the HB. Interestingly, *K. marxianus* encapsulated inside 3L-εPLL-HB reached a cell density ten times higher than growing in suspension. This could be of interest for applications in bioreactors, since it will allow to achieve higher cell densities in them or to reduce working volumes while maintaining the same microbial load. In most cases, increasing the coating from 3L to 4L had a negative effect on the cell density inside the HB. We identify two factors that can contribute to that, the lower mass transfer in 4L-HB and the toxicity of the BaCl_2_ used to crosslink the outer alginate coating layer (Figure S2). Therefore, when increasing the number of coating layers there is a trade-off between the cell density and the resistance of the HB, which can be adjusted depending on the desired application.

Microscopy observations of the εPLL-HB revealed different cellular and colony morphologies depending on the species, the number of coating layers and the culture medium. SEM images also revealed that the internal alginate 3D structure was considerably altered as a result of the microbial growth, particularly evidenced by the big internal cavities observed on 3L-εPLL-HB. Despite the variability on the external structure of the HB and the colony and cell morphology for each of the species and condition tested, we did not detect microorganisms on the surface or damage in the outer coating. This confirms that alginate core-shell HB prepared with εPLL are versatile and reliable, and can be used to encapsulate a wide variety of microorganisms with varied size, cell morphology and growing patterns.

### 4.3. Applications of εPLL-HB

Here we have demonstrated that εPLL-HB can be used for constructing spatially distributed co-cultures (SDCC) composed of species with a high disparity on their growth rates. 3L-εPLL-HB exhibited excellent reliability, enduring the mechanical stress of the vigorous shaking at 250 rpm used. More specifically, each SDCC culture consisted of twenty 3L-εPLL-HB suspended in 10 mL of medium, and no bacterial colonies were detected in any experiment after plating samples of the liquid medium. Additionally, K. marxianus growth in the SDCC was unaffected by its contact with the 3L-εPLL-HB, suggesting that the antimicrobial effect of the polymer is marginal in this co-culture configuration (Figure S1). Lastly, SDCC built with εPLL-HB offer precise control over microbial population distribution. In this study, we used a ratio of 2 HB per mL of liquid culture, but this could be adjusted to achieve any other ratio between the species in suspension and encapsulated within the SDCC. All these results confirm the robustness of the co-cultures based on εPLL-HB and lays the ground for scaling up synthetic microbial consortia using this system.

The compatibility with long-tern preservation of HB is also crucial for their applicability. The possibility to store ready-to-use HB would significantly contribute to optimize the HB manufacturing and batch-to-batch reproducibility. In this sense HB stored at −80 °C showed an optimal performance. In first place, the beads maintained their structural integrity and retained their encapsulation efficacy. Besides, *B. subtilis* viability and their growth inside 3L-εPLL-HB stored at −80 °C remained unchanged for at least 64 days of storage. To our knowledge, this is the first report on alginate core-shell beads encapsulating a bacterium that can be frozen while retaining their encapsulation efficacy unaltered. This also represent a promising result to optimize the εPLL-HB storage conditions at more moderate temperatures (Zanjani et al., 2018).

The third evaluated feature related to the applicability of 3L-εPLL-HB was their ability to protect cells against toxic and inhibitory compounds. Many lignocellulose-derived substrates are rich in phenolic compounds or furfurals produced during their pretreatments. These are toxic for some microorganisms and therefore could hinder the valorisation of these byproducts through biotechnological processes. The encapsulation inside 3L-εPLL-HB proved to be protective against the toxic effect in *P. putida* exerted by the medium M9c, derived from hydrolysed spent coffee grounds. This bacterium only grew in suspension at seeding densities above 5·10^6^ CFU/mL. However, inside 3L-εPLL-HB it survived even at the lowest seeding density tested, equivalent to 3.74·10^4^ CFU/mL. Moreover, all the *P. putida* HB showed similar cell densities, which suggests that the bacterium proliferated inside them irrespective of the number of 3L-εPLL-HB present in the well. Two protective factors could be in play. First, the higher local cell densities inside the HB could help to maintain a critical mass of viable cells. However, the M9c medium was toxic for *P. putida* growing in suspension at seeding densities up to 5·10^6^ CFU/mL. On the other hand, the local seeding cell density inside the beads is 4.5·10^6^ CFU/mL ± 1.1·10^6^ CFU/mL. This suggest that there are more effects in play that protects this bacterium against the toxic compounds other than the cell density in the local environment of the HB. More specifically, the limitations to the mass transfer of the components of the culture medium to the inside of the bead could play a beneficial role in this case, which in turn would reduce the stress over the cells and facilitate their gradual metabolization.

## 5. Conclusions

In this work we introduce an optimized method for microbial encapsulation using alginate-based core-shell hydrogel beads (HB) with ε-poly-L-lysine (εPLL) as coating agent. HB prepared with εPLL demonstrated superior microbial containment capabilities and robustness under diverse culture conditions compared to other commonly used coating materials like chitosan and α-poly-L-lysine. εPLL-HB demonstrated the ability to effectively encapsulate and at allow the growth of diverse bacterial and yeast strains with varying sizes and cell morphologies. Furthermore, εPLL-HB enabled the construction of spatially distributed co-cultures, providing an accurate control over the population balance between microorganisms with disparity in growth rates. It also proved useful for the protection against environmental stressors. Finally, they exhibited excellent performance in long-term preservation at −80 °C, maintaining their structural integrity and encapsulation efficacy, together with the microbial viability and ability to grow of the encapsulated microorganism.

Therefore, εPLL-HB offer a robust, versatile and cost-effective tool for microbial encapsulation and synthetic microbial consortia engineering, which can unlock their application in diverse biotechnological processes.

## 6. Acknowledgements

The authors acknowledge funding through grant TED2021-131194 A-I00 funded by MCIN/AEI/ /10.13039/501100011033 and NextGenerationEU/PRTR. IMR also acknowledges the financial support by grant RYC2022-037570-I funded by MICIU/AEI/10.13039/501100011033 and by ESF+.

