## Supplementary material for "Alginate-εPLL core-shell hydrogel beads as a tool for an effective, stable and scalable microbial encapsulation"

#### 1. Supplementary methods

##### 1.1. Spent coffee grounds hydrolysis

Spent coffee grounds (SCG) were provided by a local café in Granada, Spain. The lignocellulosic byproduct was firstly dried at 60 °C for 24 h. The average moisture content of raw SCG was measured using an Infrared Moisture Determination Balance (A&D Weighing, AD-4714A). Hydrothermal hydrolysis was carried out when the average moisture content of SCG was lower than 5%, as described by Perez-Burillo et al (Pérez-Burillo et al., 2019). Briefly, the reactor was filled with 10% SCG (w/v) in distilled water. The temperature was fixed to 180 °C, the stirring to 400 rpm and the reaction time to 1 h. Once the reaction finished, the reactor was quickly cooler in cold water until the temperature reached 40-50 °C. Then, the hydrolysate was filtered with a filter paper under vacuum to separate hydrochar and liquid fractions. The liquid fraction was centrifugated for 10 min at 9000 g and the pellet was discarded. Finally, the pH was adjusted to 7.3 and the hydrolysate was autoclaved at 121 °C for 30 min. Hydrolysed SCG were kept at 4 °C until use.

##### 1.2. HB preparation with chitosan as coating agent

To obtain microorganism-loaded chitosan-HB (CH-HB), a 1.5 g/L chitosan solution was prepared in a 20 mM sodium acetate and 100 mM $CaCl_2$ buffer. The mix was acidified to pH 4 with 1 M HCl, and upon dissolution of the chitosan, sterilized under UV light for 15 min. The chitosan solution was used in a 1 h incubation step for the first layer and 2 h for the second. The rest of the steps were as described in the main text.

##### 1.3. Antimicrobial assay

The antimicrobial activity of the coating polymers (chitosan, αPLL and εPLL) against each microbial strain was assessed using the agar well diffusion method (Balouiri et al., 2016). An overlay of the strain was spread onto the surface of an LB or GPY agar plate. Then, 100 µL of each coating polymer were poured into pre-made wells. The antimicrobial activity of the polymer solvents and the crosslinking solution composed of 50 mM $BaCl_2$, and 0.15 M mannitol were also tested. LB or GPY media were used as controls. Following 24 h of incubation at 30 °C, the presence of growth inhibition zones surrounding the wells was visually examined. Each condition was tested in triplicate.

##### 1.4. Stability assay of εPLL-HB in contact with suspension cultures

We tested the structural integrity of HB when incubated in contact with suspension cultures of the microbial strains. For that, a 5% (v/v) inoculum of each strain was added to a well in a 48-

well plate, with a final volume of 0.5 mL. Subsequently, four just-prepared control 3L- and 4L-εPLL-HB (not loaded with microorganism) were placed in each well. The HB were tested in M9, LB, GPY, and M9c media at 30 ºC and 300 rpm for 24 h. All conditions were run in triplicate.

### 2. Supplementary results

#### 2.1. Antimicrobial assay

Table S1. Average antimicrobial activity of the coating polymers (CH, αPLL and εPLL) against each microbial strain measured by inhibition zone diameter (mm). n.d.: indicates no inhibition zone detected.

| | Strain | | | |
|---|---|---|---|---|
| Coating polymer | ***B. subtilis*** | ***P. putida*** | ***K. marxianus*** | ***Y. lipolytica*** |
| **CH** | n.d. | n.d. | n.d. | n.d. |
| **αPLL** | 9.37 ± 0.33 | 7.86 ± 0.33 | 7.53 ± 0.24 | 6.17 ± 0.50 |
| **εPLL** | 9.99 ± 0.32 | 6.87 ± 0.14 | 7.95 ± 0.13 | n.d. |

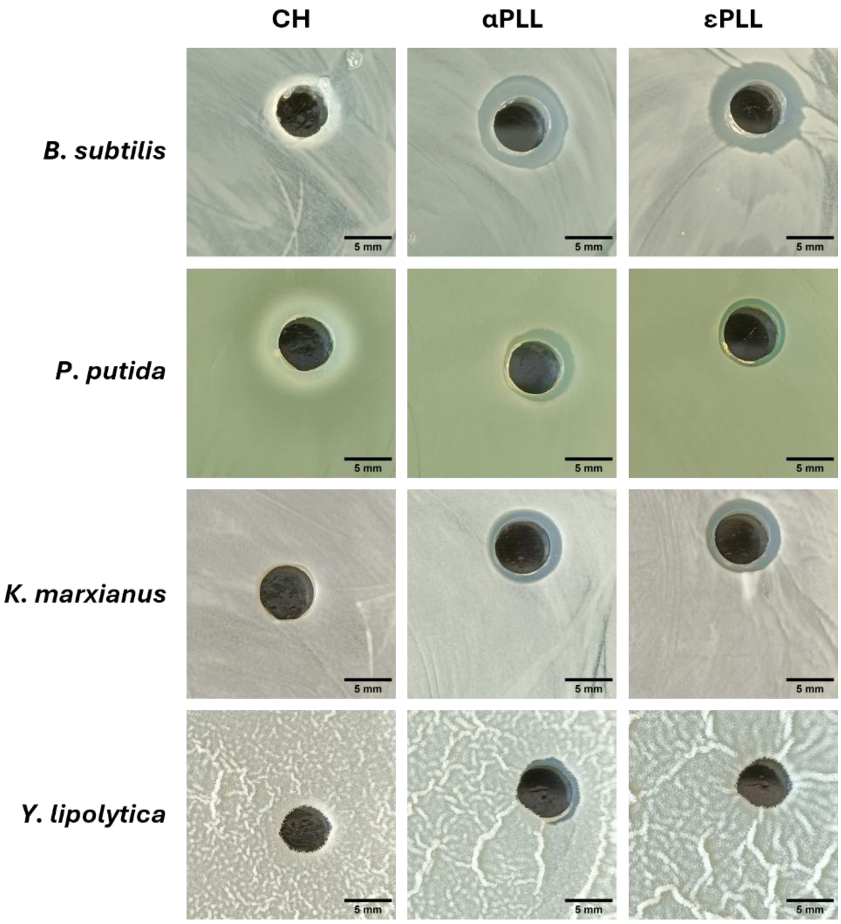


Figure S1. Antimicrobial activity of CH, αPLL and εPLL by agar well diffusion method against *B. subtilis*, *P. putida*, *K. marxianus* and *Y. lipolytica*.

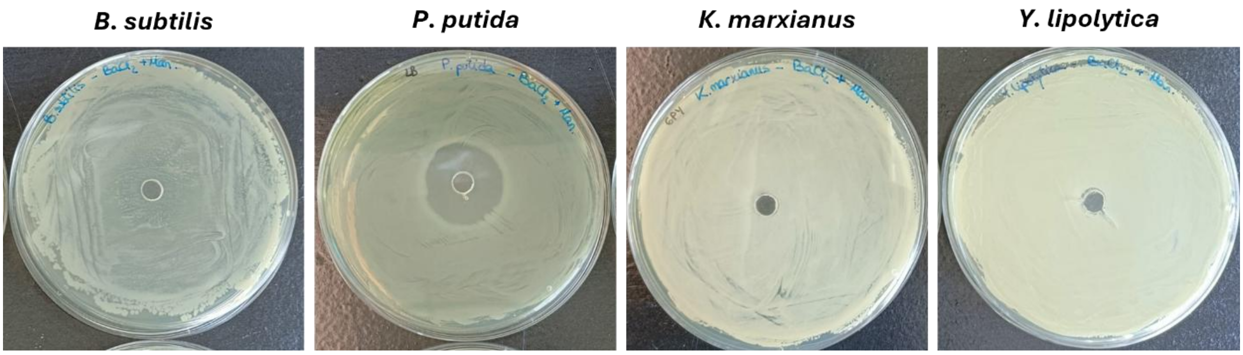


Figure S2. Antimicrobial activity of the crosslinking solution composed of 50 mM $BaCl_2$ and 0.15 M mannitol by agar well diffusion method against *B. subtilis*, *P. putida*, *K. marxianus* and *Y. lipolytica*

#### 2.2. Encapsulation efficacy of the different coating agents in rich medium

Table S2. Evaluation of the encapsulation efficacy of HB coated with CH-, αPLL-, and εPLL-HB with three (3L) or four (4L) coating layers. HB encapsulating each of the four strains tested were cultured in rich medium (LB for bacteria and GPY for yeast) for at least 48 h in individual wells. n.t: not tested

| | | Strain | | | |
|---|---|---|---|---|---|
| Coating polymer | Number of layers | ***B. subtilis*** | ***P. putida*** | ***K. marxianus*** | ***Y. lipolytica*** |
| **CH** | 3L | Escape | Escape | Escape | Escape |
| | 4L | Escape | Escape | Escape | Escape |
| **αPLL** | 3L | Escape | Escape | n.t. | n.t. |
| | 4L | Escape | Escape | n.t. | n.t. |
| **εPLL** | 3L | Containment | Containment | Containment | Containment |
| | 4L | Containment | Containment | Containment | Containment |

#### 2.3. Size of the HB core and shell

Table S3. Measurement of the alginate core diameter and the thickness (in μm) of the coating layer in 3L and 4L-εPLL-HB.

| | **HB core (μm)** | **3L-HB (μm)** | **4L-HB (μm)** |
|---|---|---|---|
| 1 | 1947 | 156 | 187 |
| 2 | 1931 | 150 | 206 |
| 3 | 1969 | 137 | 193 |
| 4 | 1903 | 138 | 190 |
| 5 | 1953 | 133 | 210 |
| 6 | 1955 | 141 | 194 |
| 7 | 1974 | 141 | 200 |
| 8 | 1957 | 144 | 187 |
| 9 | 1941 | 135 | 191 |
| 10 | 1933 | 141 | 183 |
| 11 | 1912 | 134 | |
| 12 | 1979 | 137 | |
| 13 | 1916 | | |
| 14 | 1984 | | |
| 15 | 1910 | | |
| 16 | 1943 | | |
| 17 | 1941 | | |
| **Average** | **1944** | **141** | **194** |
| **ST** | **24** | **7** | **8** |

#### 2.4. εPLL-HB size in diameter after culturing

Table S4.Diameter (in mm) of microorganism-loaded εPLL-HB after incubation in M9 and rich medium (LB or GPY for bacteria and yeast respectively). HB incubated in rich medium became wrinkled, so the measurement is approximate.

| | ***B. subtilis*** | | | | |
|---|---|---|---|---|---|
| | M9 | | | LB | |
| | 0 h | 24 h | 48 h | 24 h | 48 h |
| 3L-εPLL-HB | 1.944 | 1.896 | 1.940 | 1.889 | 1.917 |
| 4L-εPLL-HB | 1.884 | 1.931 | 1.875 | | 1.875 |
| | ***P. putida*** | | | | |
| | M9 | | | LB | |
| | 0 h | 24 h | 48 h | 24 h | 48 h |
| 3L-εPLL-HB | 1.842 | 1.784 | 1.784 | 1.789 | 1.784 |
| 4L-εPLL-HB | 1.848 | 1.872 | 1.872 | 1.870 | 1.930 |
| | ***K. marxianus*** | | | | |
| | M9 | | | GPY | |
| | 0 h | 24 h | 48 h | 24 h | 48 h |
| 3L-εPLL-HB | 1.776 | 1.809 | 1.778 | 1.671 | 1.955 |
| 4L-εPLL-HB | 1.912 | 1.865 | 1.824 | 1.453 | 1.766 |
| | ***Y. lipolytica*** | | | | |
| | M9 | | | GPY | |
| | 0 h | 24 h | 48 h | 24 h | 48 h |
| 3L-εPLL-HB | 1.752 | 1.807 | 1.768 | 1.780 | 1.702 |
| 4L-εPLL-HB | 1.847 | 1.846 | 1.743 | 1.657 | 1.659 |

#### 2.5. Characterization of microorganism-loaded εPLL-HB by microscopy

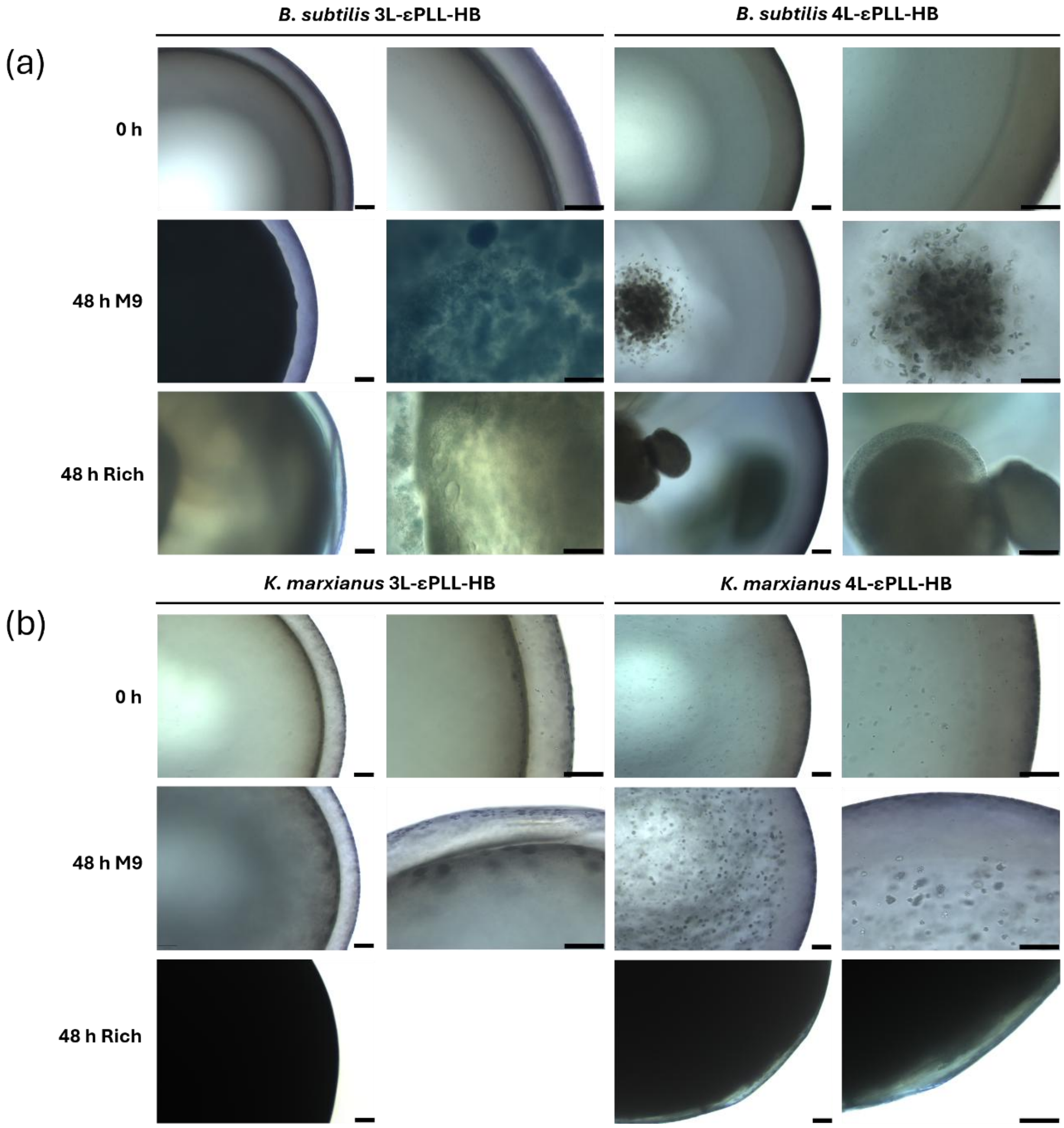


Figure S3. Microscope images of methylene blue-stained εPLL-HB with three layers (3L) and 4 layers (4L) of coating loaded with *B. subtilis* **(a)** and *K. marxianus* **(b)**. The pictures show the just-prepared HB (0 h) and those incubated 48 h in M9 or rich medium (LB and GPY for HB containing bacteria and yeast strains respectively). Scale bar: 100 µm.

Supplementary material

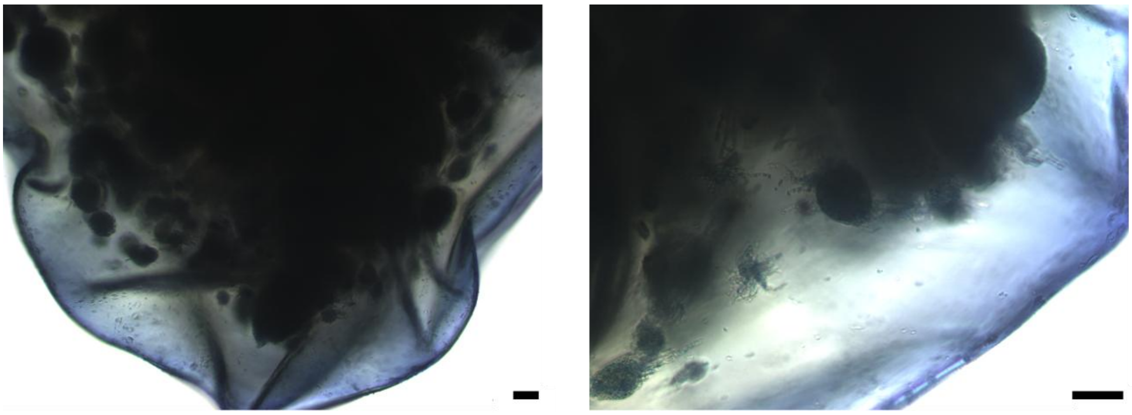

Figure S4. Microscope images of methylene blue-stained 4L-εPLL-HB containing *K. marxianus*. The pictures show the pseudohyphal morphology of *K. marxianus* after 24 h of culture in rich medium (GPY). Scale bar: 100 μm.

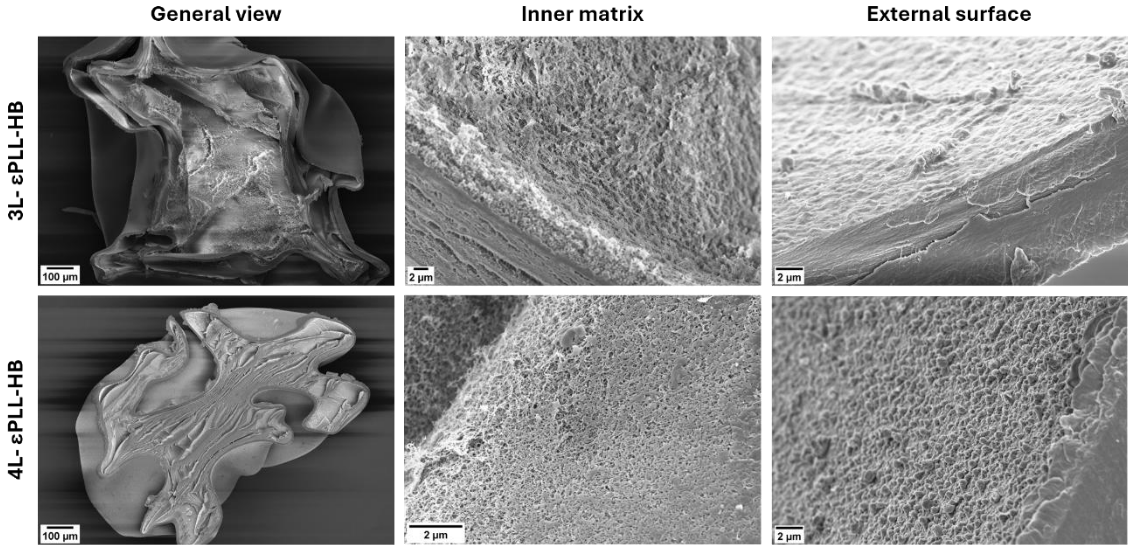


Figure S5. Cross-sectional SEM images of control εPLL-HB (not loaded with microorganism) with three layers (3L) and 4 layers (4L) of coating.

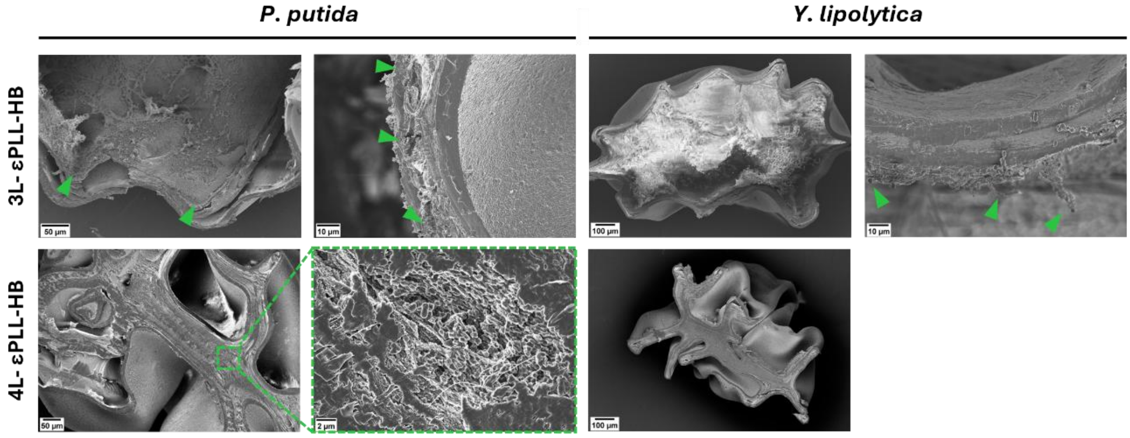


Figure S6. Cross-sectional SEM images of εPLL-HB with three layers (3L) and 4 layers (4L) of coating loaded with *P. putida* and *Y. lipolytica*. HB were incubated in M9 medium for 48 h. Green triangles indicate *P. putida* cells and *Y. lipolytica* pseudohyphae.

#### 2.6. Stability assay of εPLL-HB in contact suspension cultures

Table S5. Structural stability of control εPLL-HB (not loaded with microorganism) with suspension cultures of each strain in M9, LB, GPY and M9c media at 30 °C and 300 rpm for 24 h. n.t: not tested

| | | Strain growing in suspension | | | |
|---|---|---|---|---|---|
| Culture medium | Number of layers | ***B. subtilis*** | ***P. putida*** | ***K. marxianus*** | ***Y. lipolytica*** |
| **M9** | 3L | Stable | Stable | Stable | Stable |
| | 4L | Stable | Stable | Stable | Stable |
| **LB** | 3L | Degraded | Degraded | n.t. | n.t. |
| | 4L | Stable | Stable | n.t. | n.t. |
| **GPY** | 3L | n.t. | n.t. | Stable | Stable |
| | 4L | n.t. | n.t. | Stable | Stable |
| **M9c** | 3L | Stable | Stable | Stable | Stable |
| | 4L | Stable | Stable | Stable | Stable |

#### 2.7. Size of preserved HB

Table S6. Measurement of the diameter of 3L-εPLL-HB containing *B. subtilis* after refrigerated and frozen storage. The measures of frozen HB were carried out after defrosting and after incubation in M9 medium for 24 h.

| | **Storage** | **M9 incubation (h)** | **Diameter (mm)** |
|---|---|---|---|
| 3L-HB-1 | 4 °C | 0 | 1.835 |
| 3L-HB-2 | 4 °C | 0 | 1.896 |
| Average ± SD | 4 °C | 0 | 1.866 ± 0.031 |
| 3L-HB-1 | -80 °C | 0 | 1.796 |
| 3L-HB-2 | -80 °C | 0 | 1.730 |
| Average ± SD | -80 °C | 0 | 1.763 ± 0.033 |
| 3L-HB-1 | -80 °C | 24 | 1.898 |
| 3L-HB-2 | -80 °C | 24 | 1.734 |
| Average ± SD | -80 °C | 24 | 1.816 ± 0.082 |

#### 2.8. Protection against harsh environmental conditions

Table S7. Seeding density (CFU/mL) as a function of the number of 3L-εPLL-HB added to the 0.5 mL of M9c medium.

| nº HB | ***B. subtilis*** | ***P. putida*** | ***K. marxianus*** | ***Y. lipolytica*** |
|---|---|---|---|---|
| 1 | 9.20E+04 | 3.57E+04 | 9.03E+03 | 9.37E+03 |
| 2 | 1.84E+05 | 7.13E+04 | 1.81E+04 | 1.87E+04 |
| 3 | 2.76E+05 | 1.07E+05 | 2.71E+04 | 2.81E+04 |
| 4 | 3.68E+05 | 1.43E+05 | 3.61E+04 | 3.75E+04 |
| 5 | 4.60E+05 | 1.78E+05 | 4.52E+04 | 4.68E+04 |

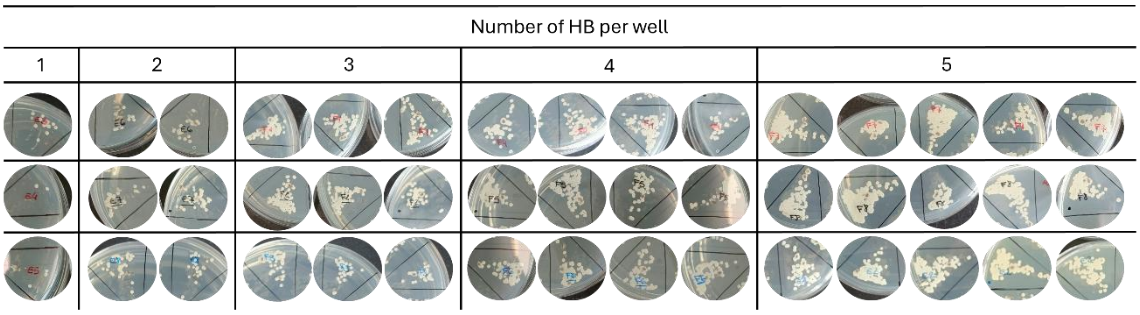


Figure S8. Colonies of *Y. lipolytica* on agar plates after the disruption of 3L-εPLL-HB incubated in M9c for 48 h. The number of HB per well ranged from one to five, and each condition was tested in triplicate. The number of colonies increased with the number of HB per well.

#### 2.9. Structural integrity of just-prepared HB and after culturing

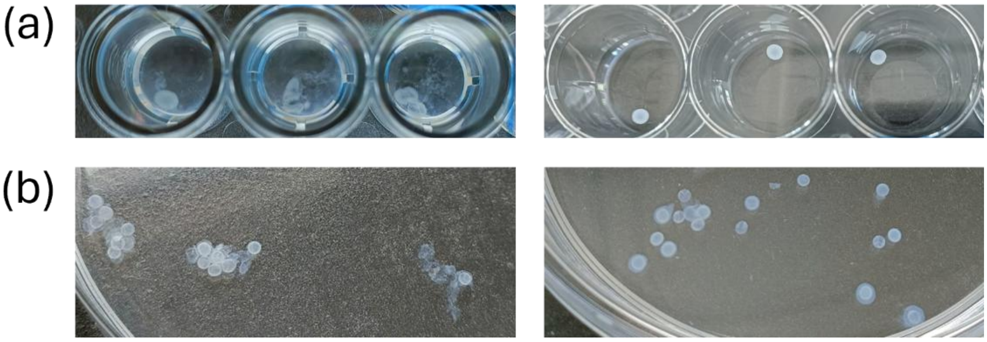


Figure S9. Effect of the chitosan coating layers on the structural stability of HB. **(a)** 3L-CH-HB lost structural integrity after 24 h of incubation in M9 medium (left), whereas 4L-CH-HB remained stable (right). **(b)** just-prepared 3L-CH-HB adhered to each other (left), while some 4L-CH-HB did not.
